# Behavioral Manipulation of *Ixodes scapularis* by *Ehrlichia muris eauclairensis*: Implications for Tick-Borne Disease Transmission

**DOI:** 10.1101/2025.03.04.641579

**Authors:** Joseph Aspinwall, Barbara Weck, Larissa A. Martins, Clayton Jarrett, Motoshi Suzuki, Mina P. Peyton, Daniel E. Sonenshine, Tais B. Saito

## Abstract

Tick-borne diseases pose significant risks to both animals and humans, with emerging pathogens like *Ehrlichia muris eauclairensis* (EME) underscoring the need for a deeper understanding of pathogen-vector interactions and tick fitness. This study investigates the impact of EME on *Ixodes scapularis* nymphs, revealing significant behavioral changes in EME-positive ticks. These ticks exhibited increased movement speed, faster bite site-seeking for attachment, and prolonged feeding durations compared to control ticks. Proteomic analyses of the tick synganglion during resting and feeding phases identified 196 differentially expressed proteins in EME-positive ticks, including multiple proteins associated with nicotinic acetylcholine signaling pathways. Our findings indicated altered neuropeptide expression related to stimulus response and activity, suggesting changes in neurophysiology. This research provides the first evidence of behavioral manipulation by an *Ehrlichia* species, indicating that the tick nervous system is a site of bacterial influence and a potential target for interventions. These findings offer new insights into pathogen-vector dynamics that could lead to the development of transmission-blocking therapies, significantly impacting tick fitness and disease transmission.

**IMPORTANCE:** Tick-borne diseases (TBDs) are increasingly affecting humans, pets, and livestock, with cases rising in recent years. Ticks can carry multiple harmful germs, and human activities and environmental are contributing to new TBDs. This study shows that the bacteria *Ehrlichia muris eauclairensis* (EME) can change the behavior of nymphal black-legged ticks, which spread various diseases. Infected ticks moved faster, attached to hosts more quickly, and fed longer than uninfected ticks. These changes were linked to specific proteins in the tick’s nervous system, suggesting that EME manipulates tick behavior. This is the first evidence that an *Ehrlichia* species can influence tick behavior, potentially increasing disease transmission. Understanding these interactions can help develop strategies to prevent TBDs by targeting the bacteria’s influence on ticks, ultimately reducing disease spread and improving public health.

## INTRODUCTION

Tick-borne diseases (TBDs) pose significant threats to human health, companion animals, and livestock, resulting in substantial economic losses (1-3). In the continental United States, TBDs account for over 75% of human clinical vector-borne disease cases (4). Since 1980, there has been a notable increase in tick-transmitted pathogens causing human diseases, with many new bacterial and viral agents identified (5, 6). Among these, several belong to the order Rickettsiales, including *Ehrlichia muris eauclairensis* (EME), which has been isolated from hundreds of clinical samples from human patients in Minnesota and Wisconsin since 2009 (7, 8).

Given the rise in TBDs, there is an urgent need to enhance our understanding of vector biology, pathogen-tick interactions, and environmental risk factors. Research into tick-pathogen interactions has revealed intricate dynamics between ticks and the organisms they carry, characterized by complex networks of symbiotic and antagonistic relationships (9-12). However, significant knowledge gaps remain, particularly regarding the mechanisms by which rickettsial pathogens influence vector transmission and the role of the tick synganglion in pathogen manipulation of tick behavior.

Studying tick-pathogen interactions is challenging due to the lack of model systems for medically significant diseases and standardized conditions for analyzing arthropod behavior (13, 14). To address this issue, our research group previously developed a tick transmission model using *Ixodes scapularis* carrying EME (15). An earlier study using a similar model found evidence of EME in the tick synganglion (16). However, the impact of these bacteria on the function of the tick’s central nervous system remains unknown. Based on these observations, we hypothesize that EME enhances the host bite site-seeking behavior and feeding efficiency of nymphal *I. scapularis* by manipulating the tick’s synganglion.

The tick synganglion has a unique structure, surrounded by a periganglionic sheath that protects the organ (17). The outer surface consists of a thin perineurium (primarily glial cells), which is believed to play an important role in the metabolic regulation of neuronal activities. Below the perineurium is the cortex or neuronal perikarya, a thicker layer containing numerous neuron cell bodies and the neuropile, an intricate layer comprising numerous axons and dendrites (17). The tick "brain" is responsible for regulating physiological processes, neurosecretory functions, and motor-associated responses (14, 18, 19). Despite the importance of these functions, little is known about interactions between *Ehrlichia* and ticks, and no studies have been conducted on how *Ehrlichia* interferes with its vector’s behavior. However, similar interactions have been demonstrated in the closely related genus *Wolbachia*, which is well-known for its complex manipulation of arthropod hosts, including behavioral alterations (20-22). Research into the impacts of *Wolbachia* on its insect hosts has led to multiple strategies for controlling mosquito populations and their pathogen transmission (23). These observations of bacteria-vector interactions in closely related genera suggest a potentially conserved mechanism within the family Anaplasmataceae.

Understanding how pathogens manipulate tick behaviors related to bite site identification, attachment, and feeding provides valuable insights into pathogen transmission (24). Further investigation into the mechanisms of behavioral manipulation could lead to novel strategies to disrupt tick bite site-seeking for attachment behavior, thereby reducing the transmission of TBDs.

This study provides new evidence concerning the relationship between EME and neuronal functions of the synganglion, which helps explain the improved fitness behavior of the EME-positive ticks. Using innovative methodologies, we demonstrate for the first time that ticks carrying EME exhibit increased speed and can locate the bite site and attach to a host’s skin significantly faster than control ticks and that behavioral changes were correlated with differential expression of neurotransmitter signaling proteins. These findings underscore the pathogen’s substantial impact on tick fitness and disease transmission. Understanding the mechanisms underlying these behavioral changes could identify potential targets for transmission-blocking therapies.

## RESULTS

In this study, we examined the effects of EME on the behavior and physiology of *I. scapularis* nymphal ticks. Given the critical importance of understanding pathogen-vector interactions due to the health risks posed by tick-borne diseases (TBDs), our research focused on the tick behavioral changes induced by EME and the potential underlying mechanisms, particularly through the manipulation of the tick synganglion. We observed significant behavioral alterations in EME-positive ticks, including increased movement speed, enhanced bite site-seeking efficiency, and extended feeding duration compared to control ticks. Furthermore, we analyzed changes protein expression in the tick synganglion to understand the neurological basis of these behavioral modifications. These findings provide new insights into pathogen-vector dynamics and suggest potential strategies for mitigating disease transmission by targeting tick behavior.

### The Presence of EME Increases Tick Attachment Rate and Feeding Time

To investigate the behavioral changes caused by EME in ticks, we examined the time required for ticks, placed in a capsule on a murine host, to migrate to the skin and attach. From the time of placement, EME-carrying ticks quickly identified a bite site and attached. The negative cohort, on the other hand, moved freely around the capsule for hours after placement. After evaluation of the attachment times, significantly more ticks from the EME-positive group were attached at both one hour and four hours after placement on a murine host (Figure 1a, b, c). When we compared the proportion of EME-positive and negative control ticks that had attached one hour after placement, 220/292 ticks from the EME-positive group were attached, compared with 133/290 negative control ticks (Figure 1a). Four hours after placement, more ticks had attached in both groups, but EME-positive ticks still attached better with 368/393 ticks feeding compared to 292/387 for the negative control cohort (Figure 1b). To account for the influence of different murine hosts, we compared the proportion of ticks that had attached per individual host. This analysis also found a significantly higher proportion of EME-positive ticks attached than negative control ticks at one hour and four hours post placement. (Figure 1c).

**Figure 1.**
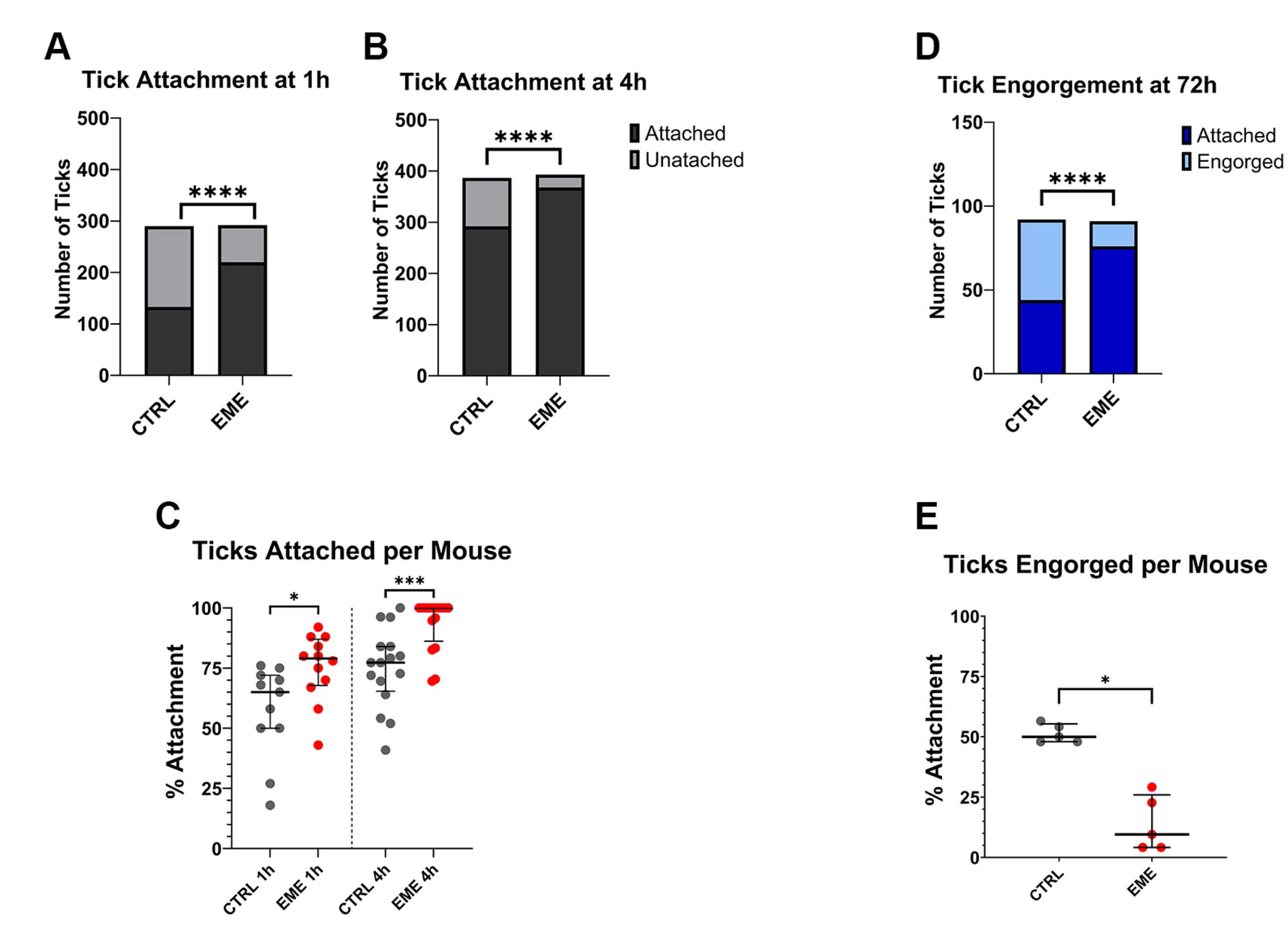
Comparison of tick attachment on the murine host and engorgement / detachment times between EME-positive (EME) and negative control (CTRL) groups. **(A)** Total number of ticks attached (light gray) and not attached (dark gray) at one-hour post-placement within the capsule on the mouse’s back. **(B)** Total number of ticks attached (light gray) and not attached (dark gray) after four hours of placement. **(C)** Scatter plot showing the proportion of ticks attached per mouse at one and four hours. **(D)** Total number of ticks still attached (dark blue) and those that had complete engorgement and detached (light blue) from the host at 72 hours post-attachment (h.p.a.). **(E)** Scatter plot showing the proportion of engorged and detached ticks per mouse at 72 h.p.a. The analysis of the number of ticks attached and detached at each timepoint was performed using Fisher’s exact test, while the number of ticks per mouse was analyzed using Mann-Whitney U test. Statistical significance is indicated as follows: (*) P-value < 0.05, (**) P value < 0.01, (***) P-value < 0.001, and (****) P-value < 0.0001. Error bars represent the median and 95% confidence intervals.

When tick engorgement and detachment at 72 h.p.a. was compared, it revealed that despite their early attachment, few EME-positive ticks from this group (15/91) had fed to repletion and detached within 72 hours compared to negative control ticks where 44/91 were engorged and detached (Figure 1d). When compared using the two strategies described above, the difference between these groups was significant independent of host influence (Figure 1e).

### The Presence of EME is Associated with Increased Tick Speed

To test motility and response to stimuli in unfed ticks, we conducted two separate experiments using tick groups placed in arenas. One arena, exposed to ambient temperature (21°C) and low humidity (40%), evaluate tick movement in response to light exposure and agitation during placement in the dish (Figure 2a, 3a, 3b). A second set of experiments under high humidity conditions (>90%) confirmed that behavioral changes were not artifacts of low humidity (Figure 2b, 3c, 3d). Under both conditions, EME-positive cohorts showed no difference in the number of ticks that moved, or the proportion of frames spent moving compared negative control ticks (Supplemental Figure 1). However, both the maximum speed and average active speed were significantly higher in EME-positive ticks in both low (Figure 4a, 4b) and high humidity experiments (Figure 4d, 4e). Interestingly, when we divided the one-hour high humidity experiment into 15 minutes sections for further comparison, a significant increase in average active speed for the EME-positive ticks compared to the negative control cohort was only observed in the first 15 minutes, while for maximum speed the increase was significant for the first 45 minutes (Figure 4c, 4f).

**Figure 2.**
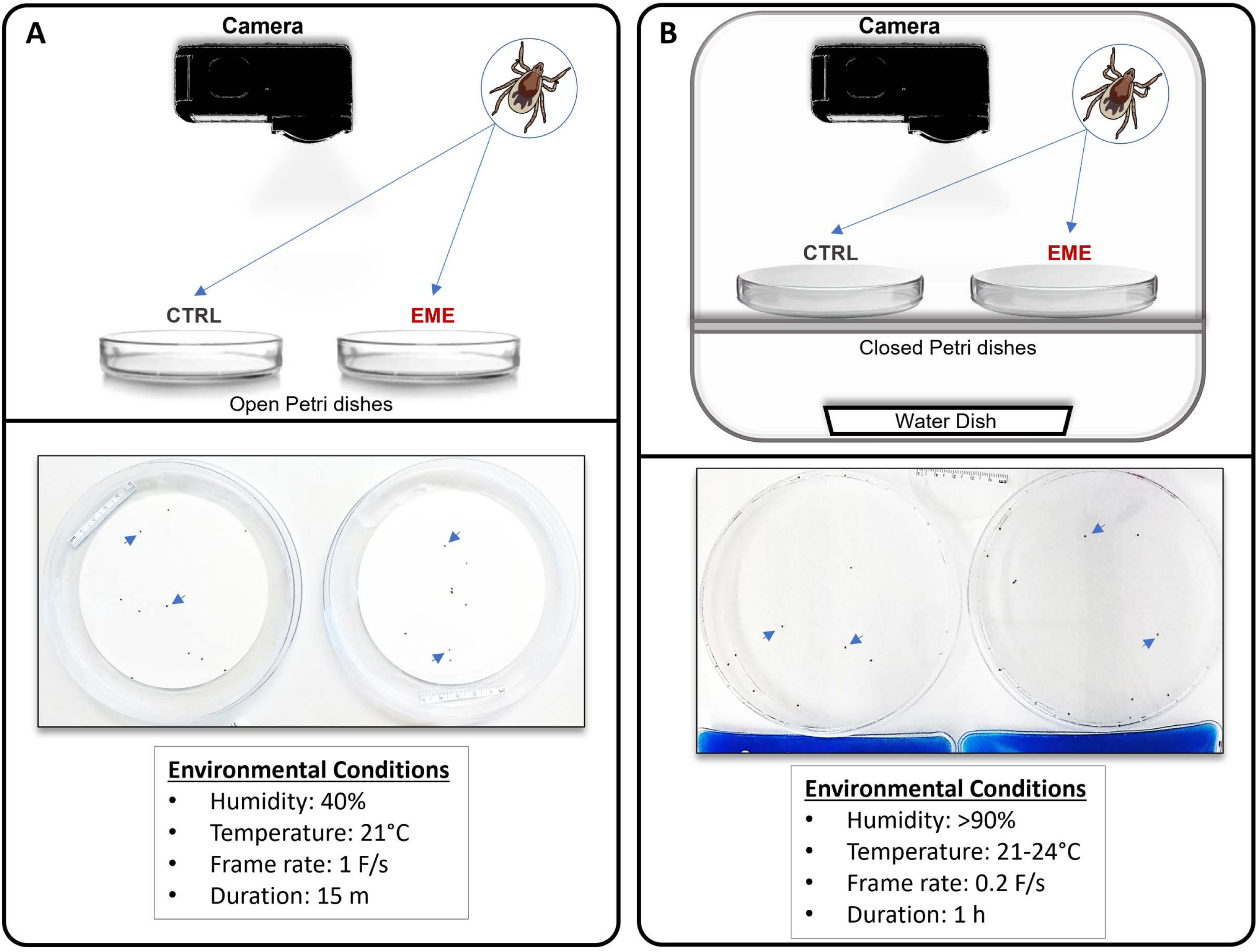
Schematic designs for evaluating off-host behavior of EME-positive (EME) and negative control (CTRL) *Ixodes scapularis* nymphs. **(A)** Experimental design for assessing tick movement under low humidity conditions. **(B)** Experimental design of the chamber used for evaluating tick movement under high humidity conditions. Experiment-specific conditions are detailed at the bottom of each panel. Blue arrows indicate examples of life ticks as captured by video camera during movement experiments.

**Figure 3.**
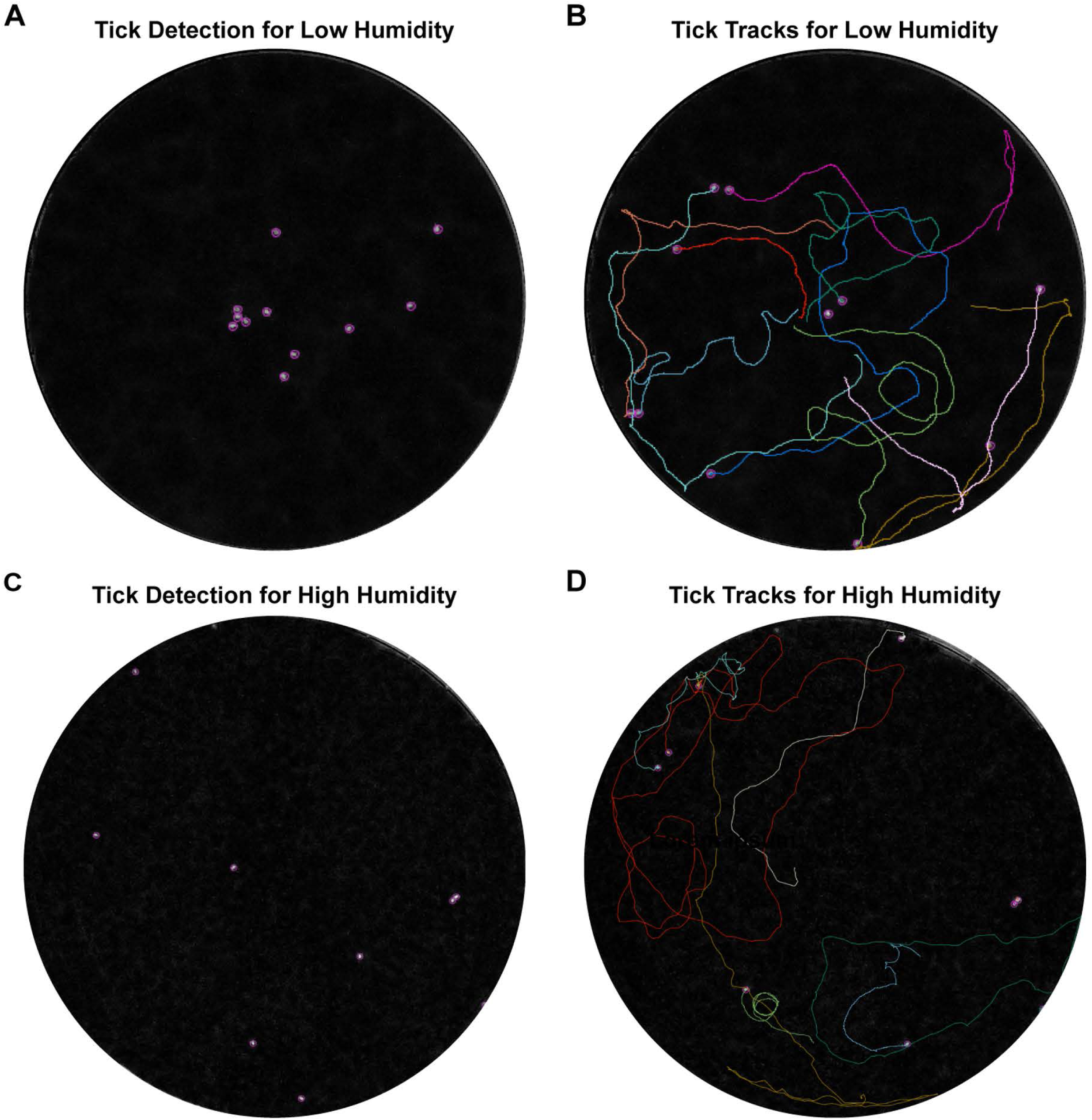
Video tracking of nymphal tick movement comparing EME-positive and negative control groups. **(A)** Tick identification in the first frame of low humidity tick movement experiment using TrackMate software. **(B)** Tick tracks overlayed on the final frame of low humidity tick movement analysis. **(C)** Tick identification in the first frame of the high humidity tick movement experiment. **(D)** Tick tracks overlayed over the final frame of the high humidity tick movement experiment.

**Figure 4.**
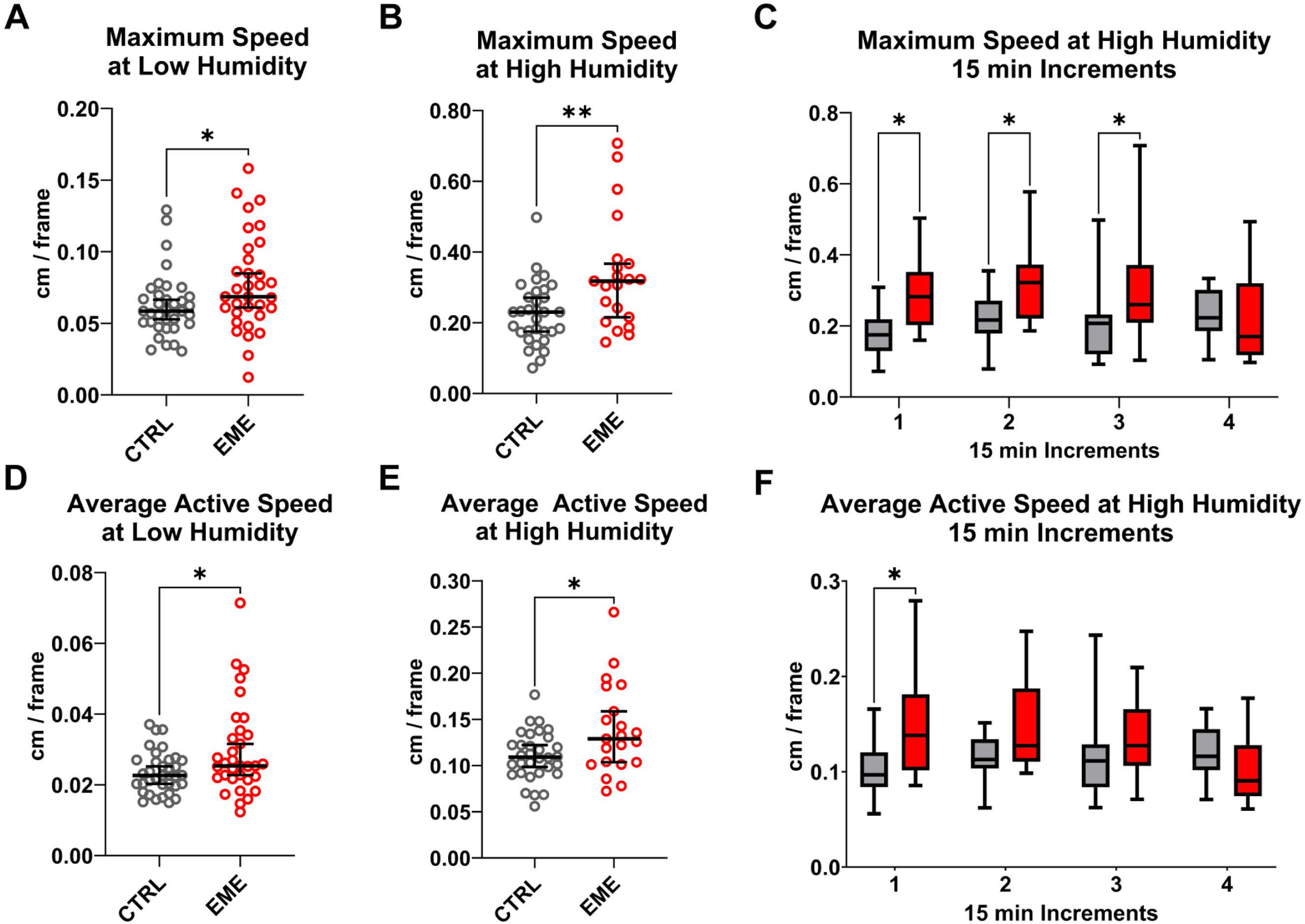
Off-host behavioral movement analysis of EME-positive (EME) and negative control (CTRL) ticks. **(A, B, C)** Measurement of maximum speed of movement between EME and CTRL nymphal ticks. **(A)** Maximum movement speed at low humidity over 15 minutes. **(B)** Maximum movement speed at high humidity over one hour. **(C)** Maximum movement speed at high humidity separated into 15-minute increments. **(D, E, F)** Measurement of average active speed of movement between EME and CTRL nymphal ticks. **(D)** Average movement speed at low humidity for 15 minutes. **(E)** Average movement speed at high humidity for one hour. **(F)** Average movement speed at high humidity separated into 15 minutes increments. All groups were analyzed using the Mann-Whitney U test. Error bars indicate the median and 95% confidence intervals. Statistical significance is indicated as follows: (*) P-value < 0.05, (**) P-value < 0.01.

### EME is Present in the Synganglion of Flat and Feeding Nymphal Ticks

At each timepoint before and during feeding, we collected the midgut, salivary glands, synganglion (Figure 5a) and remaining tissues (consolidated) to include as many tissues colonized by EME in adult ticks as possible (16). DNA quantification showed that during transmission, the tick midgut has very little bacterial presence, the salivary glands have the highest bacterial density, and the synganglion and consolidated remaining tissues both showed a similar moderate bacterial presence. With the exception of the midgut, all tissues showed increasing bacterial density during feeding (Figure 5b).

**Figure 5.**
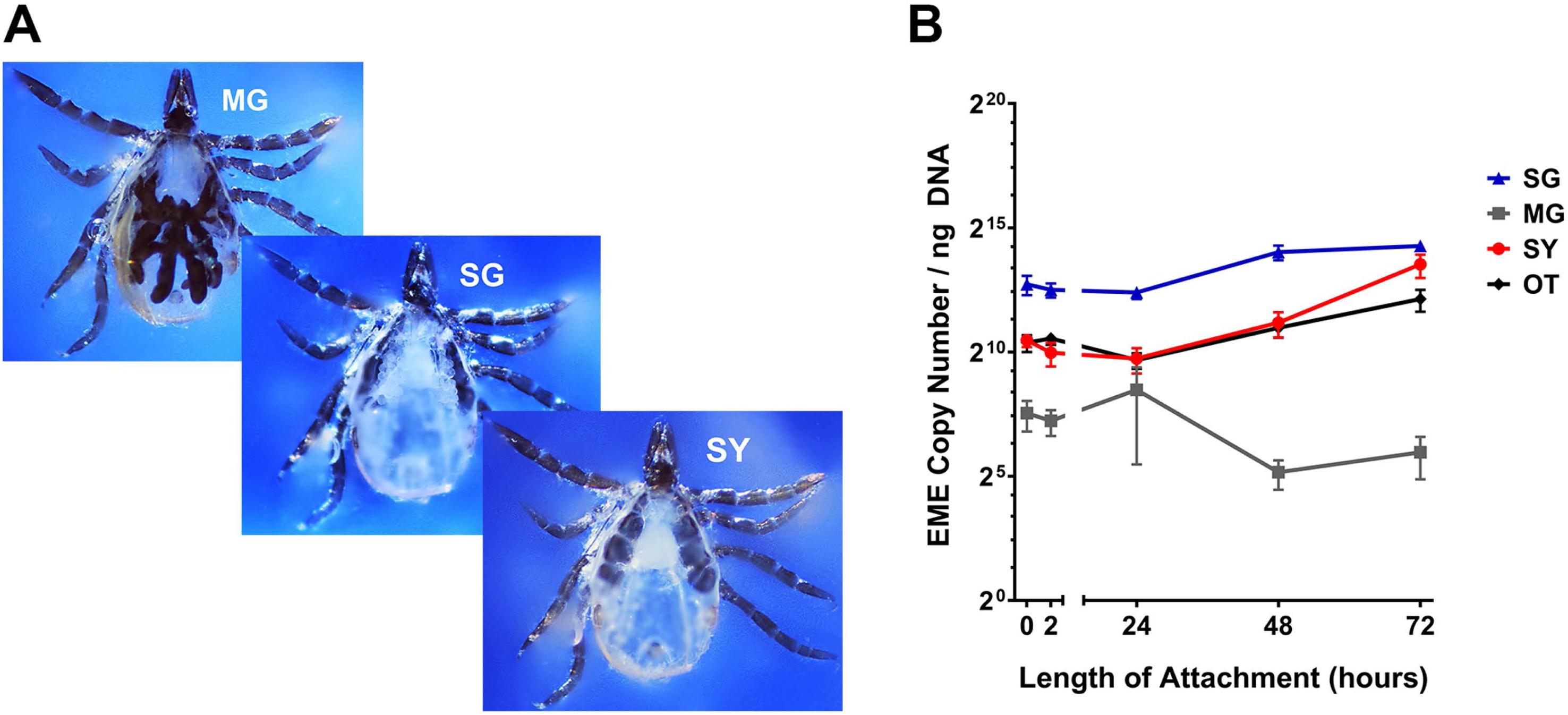
Bacterial levels in tick tissues before (0-hour) and during feeding (2 to 72 hours) on a murine host. **(A)** Representative photomicrograph of nymphal salivary glands (SG), midgut (MG), and synganglion (SY) during dissection. **(B)** Bacterial copy number per ng of DNA in different tick tissues at 0, 2, 24, 48 and 72 h.p.a. Error bars represent standard error of the mean for all samples.

### Tick *Proteins in the Synganglion*

Proteomic analysis of the synganglion from EME-positive and negative control ticks revealed the presence of mouse, tick, and bacterial proteins at all timepoints. The average total protein abundance across all timepoints was 82.6%, 17.1%, and 0.33% for tick, mouse, and bacteria, respectively (Figure 6, Figure 7). The average total number of identified proteins for tick, mouse, and bacterial proteins across all timepoints was 7137, 606, and 263 proteins, respectively. Comparing tick protein abundance, through attachment and feeding, we identify 196 tick proteins that were differentially expressed between EME-positive and negative control samples, at least at one timepoint (Supplemental Table 1). The largest number protein expression changes were observed at the 0-hour timepoint, with 42 proteins showing over-expression in EME-positive samples and 32 showing low expression (Table 1). The number of significantly affected proteins generally decreased throughout feeding, with only 18 over-expressed and 17 under-expressed in EME-positive ticks at the 72-hour timepoint. The reduced number of differentially expressed proteins over time suggests changes in tick neurophysiology and brain activity, as early time points may have a stronger stimulus-driven response that stabilizes and reaches homeostasis during feeding that lead to fewer differentially expressed proteins. Seven proteins including three uncharacterized proteins (ISCP_015976, ISCP_038280, ISCP_030944), SVEP1 (ISCP_037670), Zink metalloproteinase nas-26 (ISCP_022197), Sphingomyelin phosphodiesterase (ISCP_022951), and Cytochrome P450 3A24 (ISCP_000829) were significantly differentially expressed in EME-positive ticks across all timepoints. Of the seven, only Sphingomyelin phosphodiesterase could be linked to cellular interaction by our analysis. Protein annotation and homology revealed, protein signatures that potentially affect neurological pathways. A predicted neuropeptide, CCHamide-1 (ISCP_015817), and a protein annotated as Allatostatin A receptor (ISCP_026722) had significantly decreased expression in EME-positive ticks compared to the negative control. Additionally, a group of proteins associated with the nicotinic acetylcholine receptor signaling pathway showed differential expression between EME-positive and control ticks, indicating increased acetylcholine sensitivity (Figure 8a).

**Figure 6.**
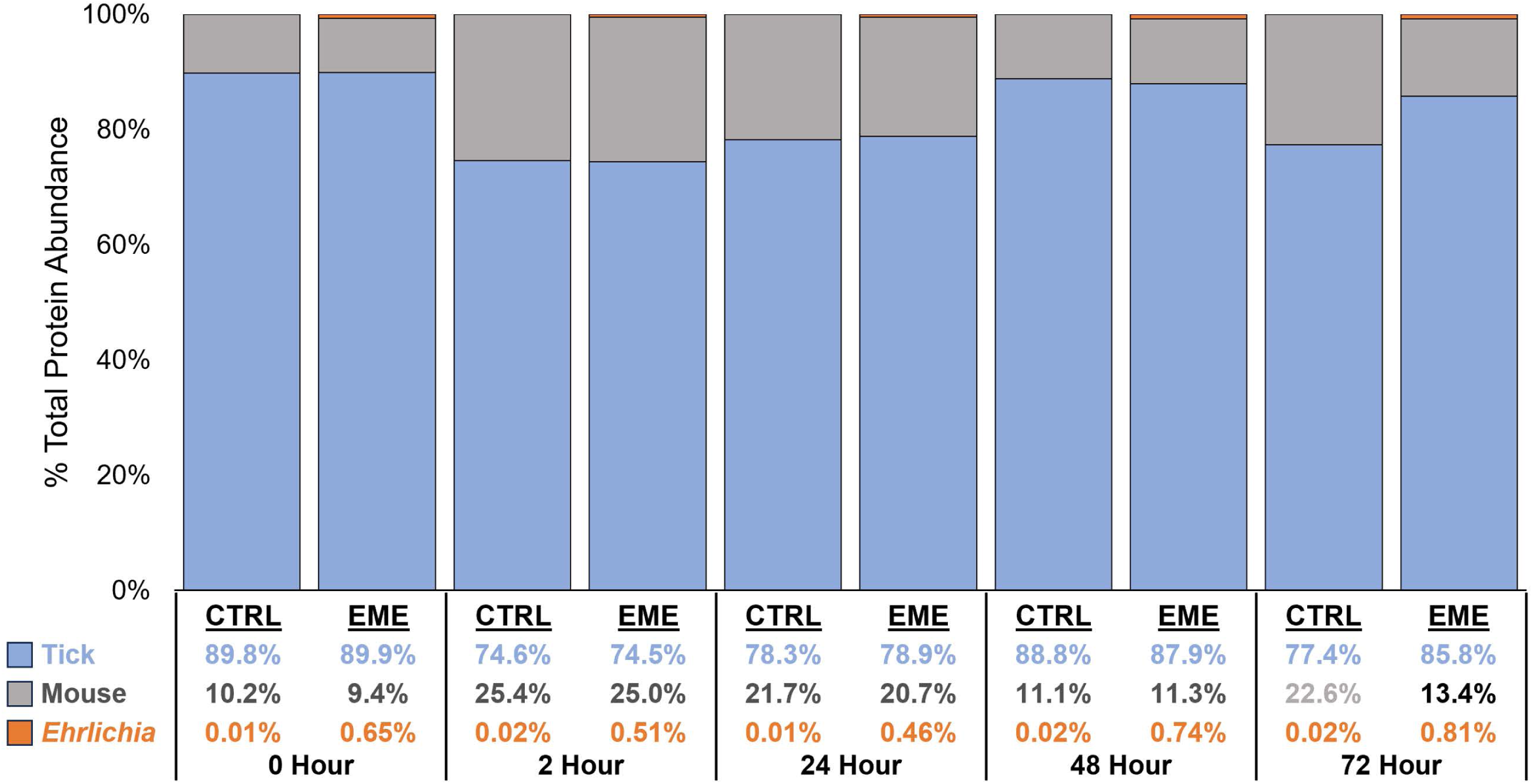
Proteomic data analysis of tick synganglion in EME-positive (EME) and negative control (CTRL) samples. Proportion of total protein abundance that are from tick, mouse or *Ehrlichia* in EME and CTRL samples across time points. Percentage total protein abundance was calculated from the average intensity values of all replicates under each condition.

**Figure 7.**
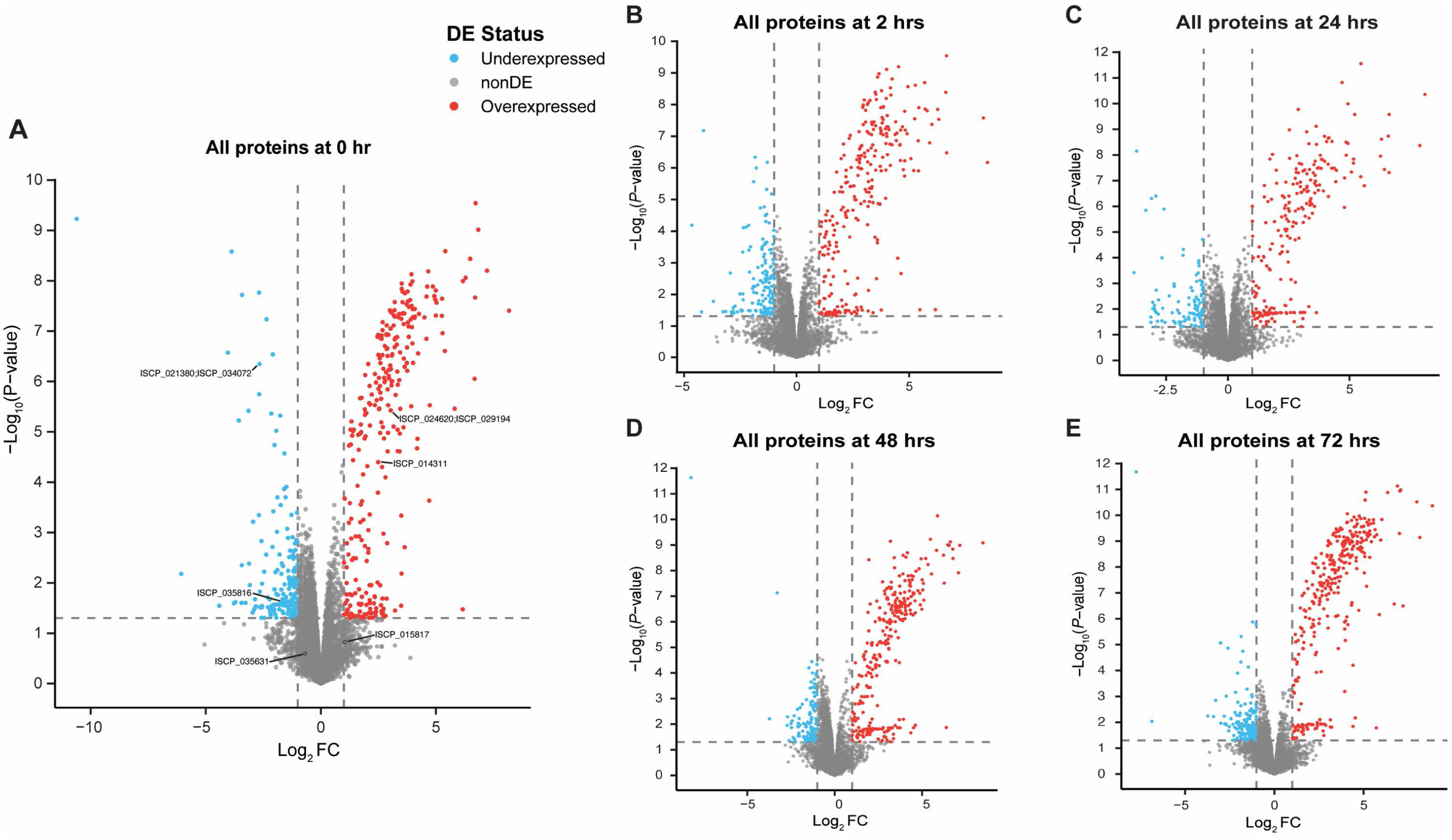
Comparative analysis of tick synganglion protein expression in EME-positive and negative control samples at different timepoints. **(A)** Volcano plot illustrating changes in protein expression at the 0-hour timepoint. Labeled proteins are associated with acetylcholine signaling or the circadian system. **(B)** Volcano plot illustrating changes in protein expression at the 2-hour timepoint. **(C)** Volcano plot illustrating changes in protein expression at the 24-hour timepoint. **(D)** Volcano plot illustrating changes in protein expression at the 48-hour timepoint. **(E)** Volcano plot illustrating changes in protein expression at the 72-hour timepoint. DE = Differentially expressed.

**Figure 8.**
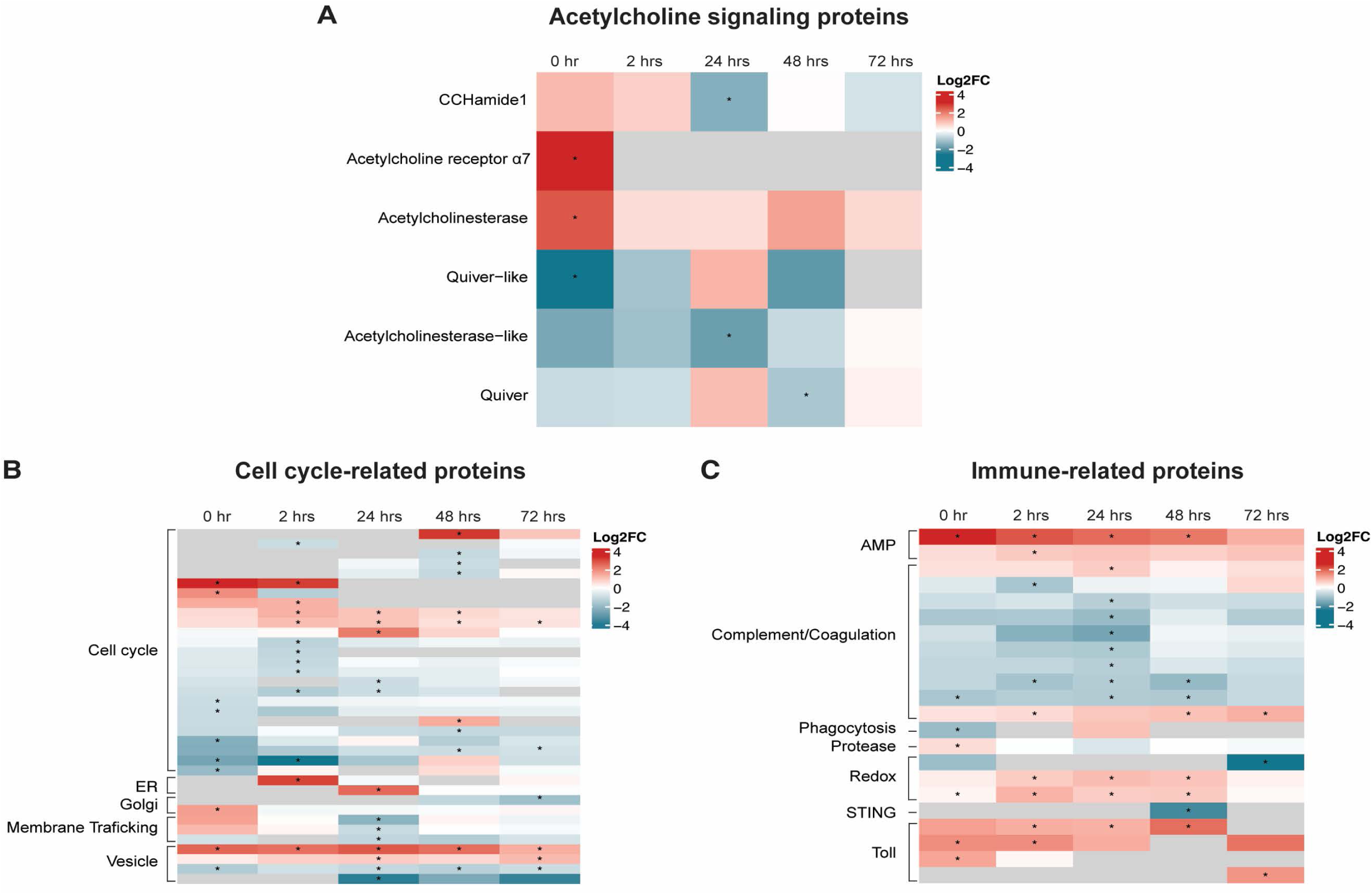
Heatmap of differentially expressed tick synganglion proteins in EME-positive group compared with negative control across all timepoints. **(A)** Proteins associated with acetylcholine signaling dysregulation. **(B)** Proteins associated with bacteria host cell interactions. **(C)** Proteins associated with the systemic immune response. Statistical significance is indicated as follows: (*) P value < 0.05.

**Table 1.** Number of tick proteins differentially expressed in the synganglion of EME-positive compared to negative control ticks at each timepoint after attachment to a murine host.

|  | 0h | 2h | 24h | 48h | 72h | Total |
| --- | --- | --- | --- | --- | --- | --- |
| High in<br>EME-positive | 42 | 41 | 30 | 20 | 18 | 84 |
| Low in<br>EME-positive | 32 | 25 | 28 | 30 | 17 | 112 |

Among the remaining protein differentially expressed between conditions, 36 were associated with the cell cycle or localization to the endoplasmic reticulum (ER), Golgi apparatus, or vacuoles (Figure 8b). VectorBase IDs and changes in expression are annotated as “cellular interaction” (Supplemental Table 1).

In addition to protein expression changes potentially linked to behavioral alterations; 22 proteins associated with innate immunity had significant changes in expression in the presence of EME. Notably, Ixoderin A (ISCP_014906) and Dae-1 (ISCP_004409) were over-expressed at multiple timepoints when EME was present in the synganglion (25, 26). Multiple Toll proteins (ISCP_016837, ISCP_024443, ISCP_012201, ISCP_015641) also showed increased expression in EME-positive ticks. Eight complement or coagulation associated proteins showed decreased expression in EME-positive ticks, however, six of these were intracellular coagulation inhibitors. P-selectin (ISCP_011173) and Complement factor B (ISCP_023386) showed increased expression (Figure 8c).

GO term analysis of proteins with differential expression identified 15/81 Wikel genome homologues to proteins with high expression in EME-positive synganglia and 42/104 for under-expressed protein. These predictions indicate a dysregulation of metabolic function and cellular processes (Figure 9a, b). When the highest E-value BLAST hits from *D. melanogaster* were used for the same analysis, 85/140 homologues were annotated, however prediction trends remained the same (Figure 9a, b).

**Figure 9.**
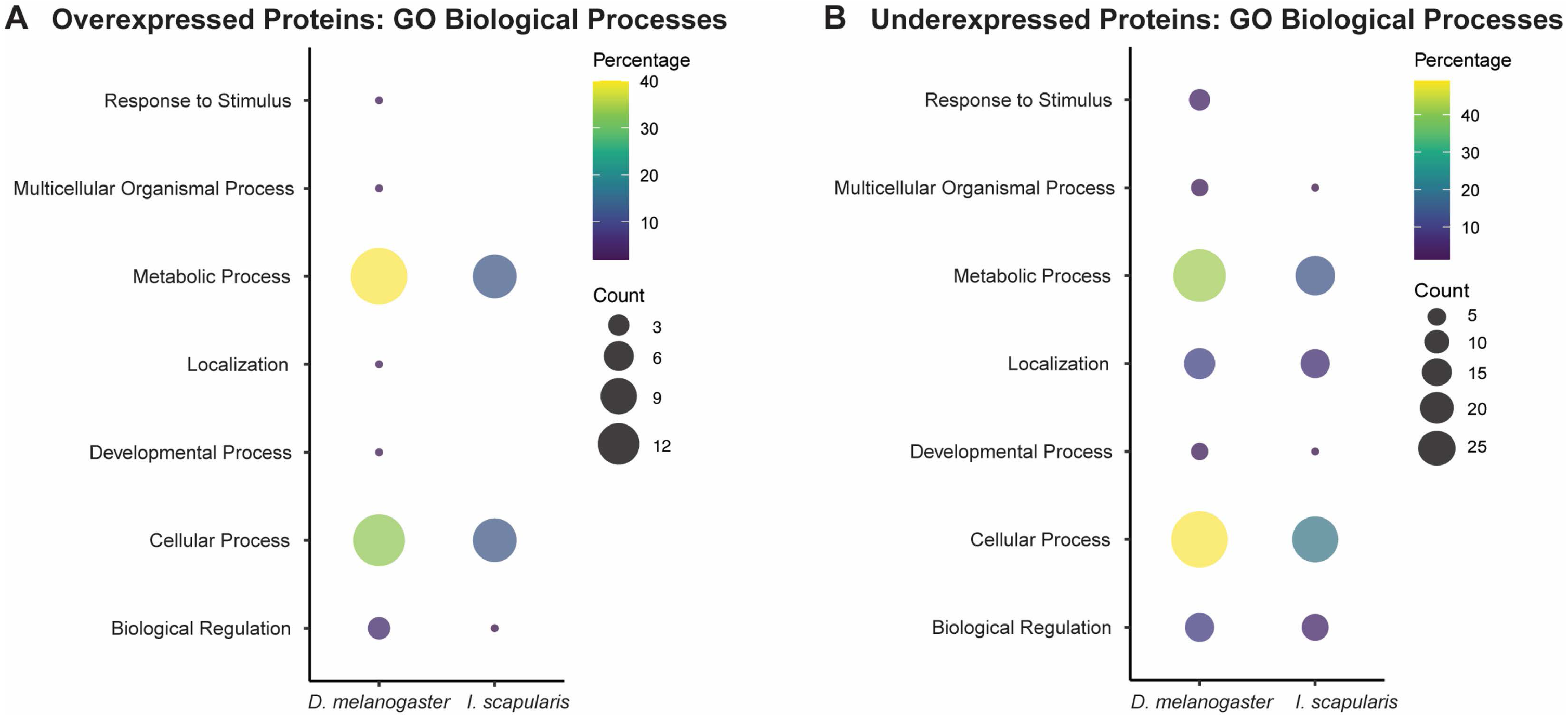
Gene ontology analysis of biological processes from differentially expressed tick synganglion proteins in EME-positive group compared to negative control at all timepoints. **(A)** Predicted GO biological processes for overexpressed proteins based on *I. scapularis* and *D. melanogaster* homologues. **(B)** Predicted GO biological processes for under expressed proteins based on *I. scapularis D. melanogaster* homologues.

### Presence of EME Proteins in the Tick Synganglion

A total of 303 EME proteins were uniquely identified in the synganglia of EME-positive ticks. Genome analysis using S4TE and Opt4e, predicted 38 of these proteins as T4SS effectors (Supplemental Table 2). Two of these (V9R5P6, V9R7B4) were homologues of ECH_0825 and ANK200, known effectors in *E. chaffeensis* (27-30). Two homologues of effectors TRP47 and TRP120 (R4QZ53, V9R5N3) secreted through the type I secretion system were also detected among bacterial proteins (31, 32).

## DISCUSSION

Our experiments demonstrated that the presence of EME in the synganglion, *I. scapularis* ticks exhibit increased movement speed, heightened aggression in seeking and attaching to bite sites, and prolonged feeding duration. Additionally, our proteomic analyses and video tracking further support the hypothesis that EME manipulates tick behavior by modulating synganglion regulation, incidentally impacting disease transmission.

When evaluating the impact of EME on nymphal behavior during infestation of a murine host, a larger proportion of EME-positive ticks successfully located and attached to bite sites faster than negative control counterparts. This suggests unique behavioral manipulation by EME leads to increased movement speed. Further, our tick movement tracking system confirms that EME-positive *I. scapularis* nymphs moved significantly faster than negative controls, independent of light and heat stimuli. The more rapidly EME-positive ticks attach to the host, less likelihood they have of being removed by the host. Therefore, our results demonstrate for the first time that ehrlichial pathogens manipulate tick vector behavior by reducing the time to seek and attach to bite sites on vertebrate hosts. Understanding the mechanisms behind these behavioral changes will help identify preventive measures against tick bites and interfere with TBD transmission. Additionally, a significantly higher proportion of EME-positive ticks remained attached for over 72 hours, indicating a longer average feeding duration. We speculate that this could enhance bacterial transmission by allowing more bacteria to be secreted into the bite site and extending the period during which uninfected co-feeding ticks could acquire EME. Further research will investigate the molecular processes that regulate increasing feeding duration.

Since the tick synganglion is responsible for neuronal regulation, including motor responses, we confirmed that EME is present in the central nervous system of nymphal *I. scapularis* before and during feeding (17). Ehrlichial presence in the cortex of the adult tick synganglion has been previously demonstrated (16), and together with our findings, suggest an effect of this bacterium on neuronal function.

To further explore the molecular mechanism by which EME interferes with the tick central nervous system, we performed proteomic studies on the synganglion of nymphal ticks during the resting phase and throughout feeding. Interestingly, our results revealed several tick-specific proteins differentially expressed in EME-positive ticks. Notably, the neuropeptide CCHamide-1 showed significant alteration in EME-positive ticks compared with control counterparts.

Studies in *D. melanogaster* demonstrated that elevated levels of CCHamide-1 are associated with delayed sleep onset and increased daylight activity (33, 34). In *I. scapularis* data indicated a diurnal sleep pattern and increased nocturnal activity, while *A. phagocytophilum* altered circadian rhythm-associated proteins in ticks (35, 36). Although experimental evidence is needed, our findings of lower CCHamide-1 levels in EME-positive ticks at 24 h.p.a. could be associated with increased feeding duration. The CCHamide-1 peptide has been also associated with starvation-related olfactory responses (37), suggesting that its differential expression could be responsible for more aggressive responses to stimuli.

EME-positive ticks also showed dysregulation in acetylcholine signaling pathways. Our results indicated that acetylcholine receptor protein expression significantly increased before tick feeding at 0 h.p.a., highlighting the importance of acetylcholine signaling in tick-EME interactions. Acetylcholinesterase was also significantly upregulated at the same timepoint, metabolizing the signal molecule and controlling cholinergic signaling. Significant amounts and widespread localization of acetylcholinesterase in the tick “brain” indicate its important role in cholinergic regulation of neuronal functions (18). If this pathway is linked to behavioral change, shortened signal duration may explain the temporary behavioral changes observed in our off-host tick movement experiments. We also found that two homologues of the *D. melanogaster* SLEEPLESS (SSS) protein, Quiver and Quiver-like, were under expressed in EME-positive ticks. In *D. melanogaster*, SSS prevents acetylcholine receptor binding and increases potassium channel protein levels, reducing signal reception and neuronal excitability (38). Deletion of SSS increases wakefulness, indicating its role in wakefulness-associated neurons.(33). A reduced expression of a tick protein like SSS in neurons expressing CCHamide-1 could increase sensitivity to acetylcholine signaling and enhance activity and wakefulness. Furthermore, increased neuronal excitability from Quiver suppression in neuronal cells’ circadian rhythms may lead to a stronger CCHamide-1 response or circadian dysregulation independent of CCHamide-1. If acetylcholinesterase upregulation occurs in the same cell as Quiver, the resulting increase in both signal sensitivity and suppression may shorten acetylcholine signaling duration.

The expression of the Allatostatin A receptor significantly decreased at 48 hours. This receptor is conserved across tick species and is associated through its binding partner with feeding regulation (39-41). Dysregulation of this pathway is likely associated with increased feeding time in EME positive ticks.

The glial cells (perineurium) surround the synganglion cortex and their cell processes extend through the nervous tissue. These cells are believed to be involved in metabolic regulation of neuronal activity (17). GO term analysis identified changes in the expression of proteins involved in metabolic activity. These included a decrease in 3-hydroxyacyl-CoA dehydrogenase levels, and an increase in histidine ammonia-lyase and imidazolonepropionase levels effecting protein and fatty acid metabolism. Interestingly, additional results showed *Ehrlichia* effectors, including mitochondrial and nuclear-targeting proteins present in the tissue, likely associated with host cell metabolism and transcription to enhance bacterial survival (27, 29, 32, 42). In glial cells, changes induced by these effectors could alter synganglion activity playing a role in EME induced changes in tick behavior.

Additional protein functions were identified for in the synganglion proteomics analysis that were not directly associated with tick behavior. They were mostly related with the tick immune response, cell processes and bacterial proteins. Across all timepoints, components of the innate immune response in EME-positive ticks showed significant changes in protein expression, including increased levels of Ixoderin A and Dae2 which effect bacterial replication, and overexpression of Toll proteins, indicating bacterial interaction and immune activation.

Additionally, EME infection has been reported to modulate cellular functions, with changes in the expression of proteins involved in vesicle transport, cell cycle regulation, and structural integrity, suggesting strategies to facilitate bacterial nutrient acquisition, replication, and survival within host cells (25, 26, 43).

In conclusion, the presence of EME in the tick vector led to significant behavioral changes during skin bite site-seeking, attachment, and engorgement. These behavioral changes were correlated with alterations in the expression of neuropeptides and neurotransmitter pathways associated with the circadian system and stimulus response in the tick synganglion. Our findings provide the first mechanistic insight into pathogen strategies for behavioral manipulation of its tick vector.

Further experimental studies are needed to precisely establish how EME in the synganglion modulates neuronal function, altering tick behavior and enhancing both tick feeding success and microbial survival. This new knowledge may also contribute to development of new methods for preventing disease transmission.

## MATERIAL AND METHODS

### Tick Acquisition of Ehrlichia

*Ixodes scapularis* ticks were obtained as eggs from Oklahoma State University (Stillwater, OK). All ticks used in our experiments were maintained in desiccators at a temperature of 21°C– 22°C, with approximately 98% humidity and darkness.

To produce an EME-positive tick cohort, we inoculated mice via tail vein with EME-infected mouse spleen homogenate, following the protocol described by Saito and Walker (15). During peak bacteremia, larval ticks were allowed to feed on infected mice. After feeding (“acquisition”), the ticks were maintained under the previously described conditions until molting. Approximately three weeks after molting, about 10% of the ticks were tested for the presence of bacteria using qPCR. Nymphal tick pools with over 90% positive for ehrlichial *dsb* gene were used for all experiments (15, 44). All mice experiments were done at NIH under the guidelines of their Institutional Animal Care and Use Committee.

### Tick Feeding and Sample Collection for Bacterial Quantification

*Ixodes scapularis* nymphal feeding followed Saito and Walker’s methods (15). Ticks were confined in a capsule on the shaved backs of C57BL/6 mice. For bacterial quantification, two replicates of ten EME-positive ticks were fed on mice for 2, 24, 48, 72 hours post attachment (h.p.a.) and unfed (0 h.p.a.). After removal from the host, ticks were washed in cold 3% bleach and 70% ethanol, rinsed in MQ-filtered water, dried on filter paper and adhered to a Petri dish. Dissections of tick midgut, salivary glands, and synganglion were performed on a 4°C cold plate, rinsed in PBS droplets, kept individually on ice, and stored at -80°C.

Tissues were suspended in 20 µL of buffer ATL (Qiagen, Germantown MD) and homogenized with a 2.8 mm stainless steel bead for two minutes at 30 oscillations/minute using the TissueLyser II (Qiagen, Germantown MD). We then followed the “mouse tail” DNA extraction protocol from the Qiagen DNeasy Blood and Tissue kit (Qiagen, Germantown MD).

Quantitative PCR used SsoAdvanced Supermix for Probes (Bio-Rad Laboratories, Hercules, CA), with primers and probe from Stevenson *et al.* (45). DNA quantification was achieved using a plasmid-generated standard curve on all PCR plates. Samples were run for 45 cycles at 60°C annealing temperature with a 30-second elongation step. Bacterial copy number was normalized to total DNA per µL, quantified using a fluorescence assay kit (Quant-it PicoGreen, Thermofisher Scientific, Waltham, MA) on Tecan Spark multiplate reader (Tecan, Switzerland).

### Tick Behavior During “Bite Site-Seeking” and Attachment, and Feeding Time

To evaluate the presence of EME within tick tissues on *I. scapularis* behaviors during skin bite site-seeking, attachment time and feeding duration, we performed a capsule-restricted infestation on naïve mice with EME-positive or negative control ticks. The same number of ticks and procedures were use for both groups. Capsules were opened, and ticks were evaluated for attachment at 30 minutes, 1, 4, 6, and 8 hours after tick placement. Consistently immobile ticks with raised abdomens were considered attached. Observations were recorded at one and four hours for 16 mice per condition and timepoint, with approximately 24 ticks per animal. At one hour, 290 negative control ticks and 292 EME-positive ticks were analyzed. Ticks were observed on 11 mice for the EME-positive cohort and 12 mice for the negative control group. At four hours, data were recorded for 16 mice in both groups, with 387 negative control ticks and 393 EME-positive ticks.

Engorgement and detachment were assessed at 72 h.p.a. for four mice per group. Replete nymphs no longer attached were recorded as engorged and detached, while attached ticks were recoded as “still attached” or feeding. Analysis included 91 ticks per group, excluding unfed ticks.

### Off-Host Tick Behavioral Experiment

To identify differences on off-host movement changes between EME-positive and control ticks, groups of ten nymphal ticks were placed on 11cm filter paper circles adhered inside Petri dishes (Figure 2a) and observed for 15 minutes. Petroleum jelly was applied around the outer edge of the filter paper. EME-positive and negative control cohorts were recorded side by side with four replicates over four days, alternating the positions of the tick arenas for each replicate. Observations were conducted at 21°C and 40%humidity (referred as “low humidity”). A second experiment was conducted in a chamber with over 90% humidity (referred as “high humidity”). Ten new ticks were placed inside a closed Petri dish allowing gas exchange with the humidified chamber. Ticks were placed in the sealed dish for over 12 hours before observation and held in a dark humidified chamber. Before observation, dishes were moved to a light-exposed chamber with a SpaceGel^TM^ chemical heat pack (Braintree Scientific, Braintree, MA) adjacent to each dish for heat stimulus (Figure 2b). Placement of EME-positive and control dishes were alternated for each replicate. Ticks were recorded for one hour after transitioning to the light-exposed chamber. This experiment had six replicates over three days, with one replicate in the morning and one in the afternoon.

### Image Capture and Behavioral Analysis

Nymphal tick movement was observed using a GoPro Hero9 camera (GoPro, San Mateo, CA) positioned above and level with the tick arenas (Figure 2). The camera’s field of view was set to narrow. Images were captured for one or five seconds for the low and high humidity experiments, respectively. The first minute of recorded frames and any additional frames after the 15 minutes or one hour observation periods were removed.

All images were imported into ImageJ and tick locations and paths were tracked (Figure 3) using the TrackMate plugin (46, 47). Thresholds were manually set to exclude false tick detections, and tracks were manually reviewed to resolve false track switching. Custom Python3 code in Jupyter Notebook version 7.0.4 was used to identify gaps in TrackMate tick detection and estimate ticks’ approximate locations and speed during the gap. Missing speeds were approximated by dividing the distance moved during the time gap by the number of frames in which the tick was undetected. Approximates coordinates were calculated by dividing the total movement in the “x” or “y” directions by the number of elapsed frames and adding these values to the previous coordinates iteratively until positions were determined for every frame within the gap.

After processing, data was analyzed for the proportion of active ticks, maximum speed, the percent of time spent moving, and the average active speed under both humidity conditions. Ticks moving slower than 0.01 cm/sec for at least one minute were classified as “still” and excluded from average active speed and maximum speed analyses. This threshold removed movement noise produce by TrackMate software. The percent of time spent moving is the proportion of frames where a tick moved more than 0.01 cm/s. The average active speed is the average distance moved by a tick in frames where it moved faster than 0.01 cm/s, and the maximum speed is the largest distance moved by a tick in a single frame. In addition to the analysis of the one-hour period for the high humidity condition, a comparative analysis was conducted in 15-minute increments for metrics showing significant differences over the entire one-hour period.

### Synganglion Collection and Extraction for Proteomics

Tick synganglion were collected from EME-positive and negative control ticks unfed and during feeding on a murine host. Four replicates of 24 unfed EME-positive and 24 unfed negative control ticks were dissected for the 0 h.p.a. timepoint. Additionally, four replicates of 24 ticks feeding per mouse, per timepoint (2, 24, 48 and 72 h.p.a.), per condition were collected. Synganglion dissections followed the method described in “Tick Dissection for Bacterial Quantification” with the addition of a protease inhibitor to the PBS. Synganglion from ticks recovered from the same mouse were pooled before extraction, with 18 to 24 synganglion consolidated per tissue pool. Samples were kept on dry ice and stored at -80°C.

For protein extraction, the synganglion samples were thawed over ice and suspended in 20 µL of RIPA buffer with 2% SDS. Samples were homogenized as described in DNA extraction, centrifuged at 16,000 rcf at 4°C for two minutes, and 30 µL of cold RIPA buffer with 2% SDS was added. After incubation at 95°C with agitation for five minutes, samples were cooled on ice and centrifuged at 16,000 rcf for eight minutes at 4°C. The supernatant was transferred to a fresh low-binding protein tube and stored at -80°C.

### Liquid Chromatography Tandem Mass Spectrometry (LC-MS/MS)

Extracted proteins were processed by the SP3 method (48). A standard solution digestion procedure was performed where extracted proteins were denatured with urea, reduced using dithiothreitol, alkylated with iodoacetamide, and digested with trypsin. LC-MS/MS data were collected using an Orbitrap Fusion Lumos mass spectrometer connected to an EASY-nLC 1200 liquid chromatography system (Thermo Fisher Scientific). The mobile phase solvent contained water and 0.1% formic acid. Approximately 900 ng of peptides mixed with iRT peptides (Byognosys) were loaded onto a trap column (PepMap 100 C18, particle size 3 μm, length 2 cm, inner diameter 75 μm, Thermo Fisher Scientific), and separated on an analytical column (PepMap 100 C18, particle size 2 μm, length 25 cm, inner diameter 75 μm, Thermo Fisher Scientific) with an acetonitrile gradient over 90 mins (2.4 - 24 % acetonitrile for 54 minutes, followed by 24 - 52.8% for 36 minutes) at the flow rate of 200 nL/min. The analytical column temperature was maintained at 50 °C. Data acquisition used the staggered data-independent acquisition method by setting 41 tMS2 scans per cycle. The isolation window was set at m/z 12 using Quadrupole. tMS2 scans for the range of m/z 394 – 886 were recorded with Orbitrap mass analyzer at the 50,000-resolution setting, and for m/z 400 – 892, with implying forbidden zones equivalent to Window Placement Optimization. The automatic gain control was set at 4e5, with a maximum injection time of 86 milliseconds. HCD fragmentation occurred at the collision energy of 25%, with the MS2 scan window set to m/z 200 – 1800. A monitor MS1 scan with Orbitrap mass analyzer at 120,000 resolution was inserted between the cycles.

### Proteomic Database Search and Analysis

Proteomic data were processed with DIA-NN (version 1.8.1) software (49). Searches were carried out against the databases of the annotated proteins from the *Ixodes scapularis* isolate PalLabHiFi (VectorBase, 7/24), mouse reference proteome UP000000589 (Uniprot, 7/24), *Ehrlichia Muris* AS145 proteome UP000018689 (Uniprot, 10/24), and common Repository of Adventitious Proteins (cRAP) from https://www.thegpm.org/ (10/24) (26, 50, 51). In addition, eight sequences of interest not found in the VectorBase database and one sequence of concatenated iRT peptides were added in the database. Precursor identification allowed for cysteine carbamidomethylation as a fixed modification, with methionine oxidation as variable modification. A 1% FDR was applied, and the “match between runs” feature was enabled. The ProteoDA R package was used to analyze proteomics results for differential expression analysis between EME-positive versus negative controls (52). DIA-NN results were filtered at 1% on the following columns: lib.qvalues, lib.pg.qvalues, quantity.quality, qvalue, pg.qvalue, and PEP in the main files. Identified proteins were also filtered for minimum replicates per group, n – 1, where *n* is the number of samples per group and only proteins with minimum replicates per group were kept for further analysis. Proteins with missing values was imputed using 1% of the minimum value detected. Quantile normalization and log_2_ transformation was then applied. Proteins were considered significant and differentially expressed if they had an adjusted p-value < 0.05 and log_2_ fold-change ≥ |1|.

Tick proteins differentially expressed between conditions were analyzed using a selected list of potential neuropeptides and receptors and immunity-related proteins from *Ixodidae* ticks literature (25, 39, 41, 53-57). Proteins identified in other genomes were linked to the best protein BLAST hit from the PalLabHiFi genome using BLAST+ version 2.16.9 based on E-values and hit length (58). For neuropeptides without a BLAST hit, homologues from a closely related *Ixodes* genome were added to the search database for DIA analysis. OrthoMCL gene groups were used to identify orthologue groups (50). Affected protein sequences were queried against *Drosophila melanogaster* proteins from FlyBase (59), with hits below an E-value 10^-10^ used to interrogate protein function. Proteins were assessed for gene ontology (GO) terms using PANTHER tools (60). Reciprocal best BLAST hits from the *I. scapularis* Wikel proteins (GCA_000208615.1) against the PalLabHiFi genome and *D. melanogaster* hits were used for GO term biological process analysis. Experimentally validated effector genes from *E. chaffeensis* were taken from the literature and used to identify homologues in the *E. muris* AS145 genome by protein BLAST comparison (27, 29, 31, 32, 42). Type IV effectors prediction tools S4TE and OPT4E identified additional proteins of interest (28, 30).

### Statistical Analysis

Tick movement, attachment and detachment was analyzed using GraphPad Prism version 10.1.2 for Windows (GraphPad Software, Boston, MA). A two-way ANOVA analysis followed by a Tukey’s range test was used for comparison of bacterial density in different tick tissues. Fisher’s exact test assessed tick attachment, detachment, and the proportion of moving ticks. Mann-Whitney tests evaluated tick average and maximum speed, time spent moving, and the proportion of ticks attached and engorged/detached per mouse. Multiple Mann-Whitney tests with Holm-Šídák correction were used to compare average and maximum speed over 15 minutes increments. Differential expression analysis was performed using the ProteoDA package, which imports functions from limma for statistical analysis, applying Benjamini-Hochberg method for multiple comparisons.

## ACKNOWLEDGEMENTS

The authors extend their gratitude to Glenn Nardone of the Proteins & Chemistry Section at the Research Technologies Branch (RTB), National Institute of Allergy and Infectious Diseases (NIAID), NIH, for his experience and technical support with mass spectrometry data acquisition. We also acknowledge Roky Mountain Veterinary Branch (RMVB) for their assistance with animal care. Additional thanks to the Visual Arts Section of the RTB, NIAID, NIH, for their help with figures.

## Funding

This work was funded by the Intramural Research Program of the Division of Intramural Research (Project AI001344-04), and in part by the BCBB Support Services Contract HHSN316201300006W/75N93022F00001 to GUIDEHOUSE, INC., both form the National Institute of Allergy and Infectious Diseases (NIAID), National Institutes of Health, Department of Health and Human Services.

## Author contributions

Study designed and conceptualization: TBS, JA, DES; Experiments execution and methodology: TBS, JA, BW, LAM, CJ, MS; Data analysis: JA, MS, MPP, TBS, DES; Manuscript preparation: JA, TBS, DES, MS, MPP, LAM, CJ, BW; Funding acquisition: TBS, MPP; Supervision: TBS.

## Competing interests

All other authors declare they have no competing interests.

## Data and materials availability

The mass spectrometry proteomics data have been deposited to the ProteomeXchange Consortium via the PRIDE (61) partner repository with the dataset identifier PXD060563. Code used for movement analysis can be accessed by request to the author. All other data is available in the text or supplementary materials.

## SUPPLEMENTAL MATERIAL

**Supplemental Figure 1.** Tick movement analysis during off-host behavioral experiments comparing EME-positive (EME) and negative control (CTRL) ticks. **(A)** Total number of recorded ticks under low humidity condition, categorized as “Still” (light gray) and “Moving” (dark gray). **(B)** Scatter plots showing the proportion of recorded frames, under low humidity conditions, in which a tick was identified as moving. **(C)** Total number of recorded ticks under high humidity conditions, categorized as “Still” (light gray) and “Moving” (dark gray). **(D)** Scatter plots showing the proportion of recorded frames, under high humidity conditions, in which a tick was identified as moving. Statistical analysis was conducted using Fisher’s Exact test or Mann-Whitney U test. Error bars represent the median and 95% confidence interval.

**Supplemental Table 1.** Tick proteins with significantly different expression in EME-positive tick synganglion compared to negative control ticks at least one timepoint before (0 hour) or after (2 to 72 hour) attachment to a murine host

**Supplemental Table 2.** EME proteins identified in negative control tick synganglion in any of the proteomic analyses conducted before or during feeding on a murine host

**Supplemental Table 3.** Results from two-way ANOVA analysis and Tukey’s tests of EME density in nymphal *Ixodes scapularis* tick tissues before and during feeding on a vertebrate host

